# Development of a highly sensitive luciferase-based reporter system to study two-step protein secretion in cyanobacteria

**DOI:** 10.1101/2021.10.08.463751

**Authors:** David A. Russo, Julie A. Z. Zedler, Fabian D. Conradi, Nils Schuergers, Poul Erik Jensen, Conrad W. Mullineaux, Annegret Wilde, Georg Pohnert

## Abstract

Cyanobacteria, ubiquitous oxygenic photosynthetic bacteria, interact with the environment and their surrounding microbiome through the secretion of a variety of small molecules and proteins. The release of these compounds is mediated by sophisticated multi-protein complexes, also known as secretion systems. Genomic analyses indicate that protein and metabolite secretion systems are widely found in cyanobacteria; however little is known regarding their function, regulation and secreted effectors. One such system, the type IVa pilus system (T4aPS), is responsible for the assembly of dynamic cell surface appendages, type IVa pili (T4aP), that mediate ecologically relevant processes such as phototactic motility, natural competence and adhesion. Several studies have suggested that the T4aPS can also act as a two-step protein secretion system in cyanobacteria akin to the homologous type II secretion system in heterotrophic bacteria. To determine whether the T4aP are involved in two-step secretion of non-pilin proteins, we developed a NanoLuc-based quantitative secretion reporter for the model cyanobacterium *Synechocystis* sp. PCC 6803. The NLuc reporter presented a wide dynamic range with at least one order of magnitude more sensitivity than traditional immunoblotting. Application of the reporter to a collection of *Synechocystis* T4aPS mutants demonstrated that two-step protein secretion in cyanobacteria is independent of T4aP. In addition, our data suggest that secretion differences typically observed in T4aPS mutants are likely due to a disruption of cell envelope homeostasis. This study opens the door to explore protein secretion in cyanobacteria further.

**Importance:** Protein secretion allows bacteria to interact and communicate with the external environment. Secretion is also biotechnologically relevant, where it is often beneficial to target proteins to the extracellular space. Due to a shortage of quantitative assays, many aspects of protein secretion are not understood. Here we introduce a NanoLuc (NLuc)-based secretion reporter in cyanobacteria. NLuc is highly sensitive and can be assayed rapidly and in small volumes. The NLuc reporter allowed us to clarify the role of type IVa pili in protein secretion and identify mutations that increase secretion yield. This study expands our knowledge on cyanobacterial secretion and offers a valuable tool for future studies of protein secretion systems in cyanobacteria.

## Introduction

Cyanobacteria are metabolically versatile microbes that can be found in virtually every terrestrial and aquatic habitat. Their wide distribution is due to the ability to respond rapidly to environmental fluctuations and establish new niches. One of the major strategies through which cyanobacteria respond to environmental stimuli is the production and release of a large repertoire of proteins and metabolites (1–3). Protein translocation across the outer membrane into the extracellular space is known as secretion and is typically mediated by highly specialized multi-component protein complexes known as secretion systems. In Gram-negative bacteria, at least ten protein secretion systems have been identified (4). However, only four have been found in cyanobacteria: the type I secretion system (T1SS), type IV secretion system (T4SS), type V secretion system (T5SS) and the type IVa pilus system (T4aPS) (5). The T1SS and the T4SS are one-step secretion systems that connect the cytoplasm directly to the extracellular environment, thus bypassing the periplasmic space. The T5SS and T4aPS are two-step secretion systems that rely on the general secretory (Sec) and the twin-arginine translocation (Tat) pathways for protein export to the periplasm before mediating secretion across the outer membrane (5). In addition, secretion can also occur through unconventional pathways, such as the release of extracellular vesicles (6). Overall, cyanobacterial protein secretion systems remain poorly described and little is known regarding their distribution, function and regulation.

In recent years, the T4aPS from cyanobacteria have been characterized in some detail. Type IVa pili (T4aP) are filamentous surface appendages involved in a range of processes including phototactic motility, natural competence, aggregation and flotation (7–10). The T4aPS, or elements thereof, can be found in virtually all cyanobacteria, including those where natural competence or motility has not yet been described (5). The assembly of T4aP on the cell surface is the result of the coordination of multiple proteins to form an intricate nanomachine. This complex is anchored in the cytoplasmic membrane and extends across the periplasm and outer membrane to the extracellular space. The cyanobacterial T4aPS consists of several highly conserved core units: a retractable pilus fiber, mainly constituted of the major pilin PilA, an assembly platform (PilC), ATPases that extend (PilB) and retract (PilT) the pilus fiber, an outer membrane pore through which the pilus fiber extends (PilQ), a bifunctional peptidase/methylase (PilD) that cleaves the signal peptide of nascent pilins and a complex that aligns the pilus fiber through the periplasm (PilMNOP). Cyanobacteria also encode several minor pilins that play a role in natural transformation and aggregation and may be incorporated as components of the pilus fiber (11, 12).

The T4aPS is evolutionarily, structurally and functionally similar to bacterial type II secretion systems (T2SS) (13, 14). The major differences are that the T2SS lacks a retraction ATPase and has a shorter pilus fiber, termed pseudopilus, which extrudes folded proteins across the outer membrane (15). Typically, the T4aPS is exclusively responsible for the biogenesis and assembly of pilin proteins. However, in some bacterial genomes the secretion of some non-pilin proteins depends on the correct functioning of the T4aPS (16–19). This has led to the notion that the T2SS and the T4aPS may share subunits of their machinery during the two-step secretion of non-pilin proteins (16, 20, 21). Cyanobacteria lack a canonical T2SS, however recent studies have shown that the disruption of T4aP assembly leads to dysregulated levels of secreted proteins (22, 23). Despite these observations, it remains unclear whether the cyanobacterial T4aPS is directly involved in the transport of non-pilin proteins from the periplasm to the extracellular space.

Previously we have shown that the fast-growing cyanobacterium *Synechococcus elongatus* UTEX 2973 can secrete the heterologously expressed enzyme *Tf*AA10A, a lytic polysaccharide monooxygenase (LPMO) from the gram-positive bacterium *Thermobifida fusca. Tf*AA10A was secreted via a two-step process with the cleavage of its N-terminal Sec signal peptide occurring when crossing the cytoplasmic membrane (24). To address whether the T4aPS can secrete non-pilin proteins, we developed a NanoLuc luciferase (NLuc)-based secretion reporter harboring the *Tf*AA10A signal peptide and genetically transferred the reporter into a collection of cyanobacterial T4aPS mutants. We show that the T4aPS is not directly involved in the secretion of non-pilin proteins. Instead, the changes to the exoproteome previously observed in T4aPS mutants may be due to pleiotropic effects.

## Results

### NLuc is efficiently secreted in Synechocystis sp. PCC 6803

In our previous study, we demonstrated the secretion of the LPMO *Tf*AA10A from *T. fusca* in the cyanobacterium *S. elongatus* UTEX 2973 (24). *Tf*AA10A is a copper-containing enzyme with a complex catalytic cycle not suitable for high-throughput analysis. To quantitatively test protein secretion levels in a high-throughput manner, the N-terminal Sec signal peptide from *Tf*AA10A was fused to NanoLuc luciferase (NLuc). NLuc is a 19.1 kDa luciferase enzyme from the deep-sea shrimp *Oplophorus gracilirostris* that converts the substrate furimazine to produce high-intensity luminescence (25). The *Tf*AA10Asp-NLuc fusion was inserted into the multiple cloning site of the self-replicating broad-host-range pDF-trc expression vector (26) to generate the plasmid pDAR25 (Fig. 1A) and conjugated into the motile ‘PCC-M’ strain of *Synechocystis*. We chose the ‘PCC-M’ *Synechocystis* strain for this study due to the extensive knowledge on its T4aPS and the existence of several well-characterized T4aPS mutants (9, 27, 28).

**Figure 1.**
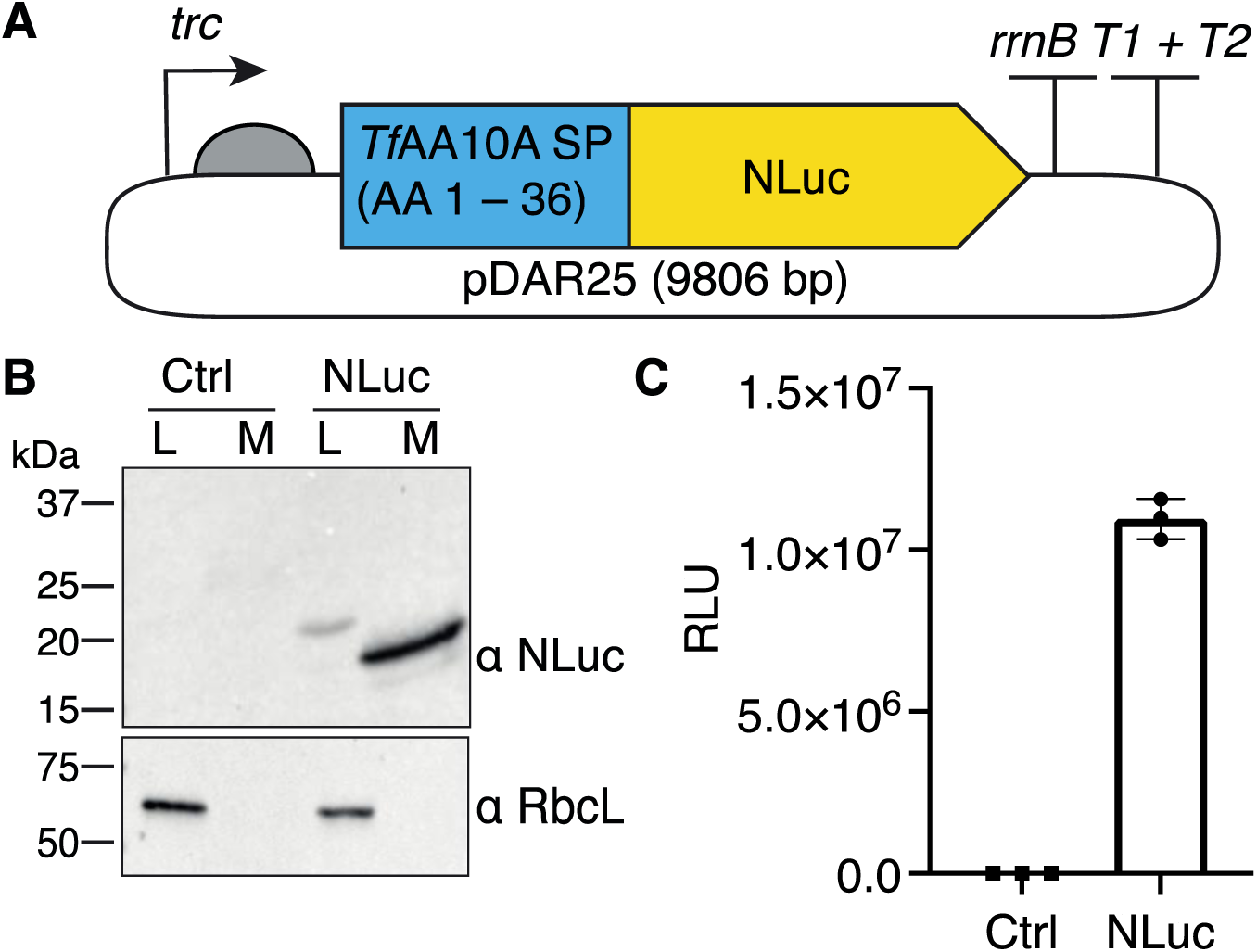
Overview of *Tf*AA10Asp-NLuc expression in *Synechocystis*. A) *Tf*AA10Asp-NLuc encodes the native protein sequence of NanoLuc carrying the N-terminal signal peptide from *Tf*AA10A. B) Detection of NLuc in the cellular lysate (L) and culture medium (M) by immunoblot analysis using an NLuc antibody (α NLuc). Ctrl designates a pDF-trc empty vector control strain. An antibody against RbcL (α RbcL) was used to verify that the NLuc in the medium did not derive from cell lysis. Lanes are normalized to 10 µg of total protein. C) Detection of NLuc in the medium by luminometry. Lanes are normalized to 1 µg of total protein.

To confirm expression of NLuc, mid-exponential cultures of *Synechocystis*, harboring either the pDAR25 or the empty pDF-trc control plasmid, were diluted to OD_750 nm_ = 0.5 and induced by the addition of 100 µM Isopropyl β-D-1-thiogalactopyranoside (IPTG) for 48 h. The cleared cellular lysate and cell-free culture medium of both strains were then separately analyzed with an anti-NLuc antibody (Fig. 1B). In the cellular lysates of the NLuc strain, only a faint band could be detected at a molecular weight matching the unprocessed *Tf*AA10Asp-NLuc fusion (approximately 23 kDa). In the culture medium, only the mature NLuc (approximately 19 kDa) was found. As expected, in the empty vector strain no bands were detected in the cellular lysate or the culture medium. To confirm that the presence of the protein in the extracellular space was due to secretion rather than cell lysis, both culture medium and cellular lysate samples were analyzed for the presence of the large subunit of ribulose-1,5-bisphosphate carboxylase/oxygenase (RbcL). RbcL could only be detected in the cellular lysates (Fig. 1B). Finally, to determine if the protein was correctly folded and active, we proceeded to assay 1 µg of total protein from the culture medium by luminometry. Luminescence could be detected in the culture medium of the *Synechocystis* NLuc strain with a negligible background signal observed in the empty vector control (Fig. 1C). Overall, these results indicate that the *Tf*AA10A signal peptide can efficiently direct NLuc, in an active state, to the extracellular space of *Synechocystis*.

### NLuc is a sensitive secretion reporter with a wide dynamic range suitable for high-throughput screening in cyanobacteria

With the confirmation that *Synechocystis* secretes NLuc we proceeded to test its application as a secretion reporter. Assaying secretion by immunoblot analysis is technically challenging due to low sensitivity and long processing times. Therefore, we set out to determine if we could replace this tedious procedure with simple luminescence measurements. We started by comparing NLuc performance by immunoblotting and luminometry (Fig. 2A, B). First, a serial dilution of culture medium was made to prepare immunoblot samples ranging from 1 µg to 31.25 ng of total protein. Equivalent samples were also prepared for luminometry. The NLuc signal detected by immunoblotting decreased linearly and was visible down to 125 ng. By luminometry the NLuc signal decreased linearly down to 31.25 ng, equivalent to approximately 1.5 µL of culture medium in our assay, where it registered an average value of 287,500 ± 5,700 RLU. Considering the low amount of background luminescence observed in the control reactions (Fig. 1C), the NLuc assay has the potential to be scaled down to detect sub-nanogram amount of secreted protein in sub-microliter volumes. These experiments demonstrate that, when compared to immunoblotting, NLuc has higher sensitivity, wider dynamic range, and can detect secreted proteins in minimal volumes.

**Figure 2.**
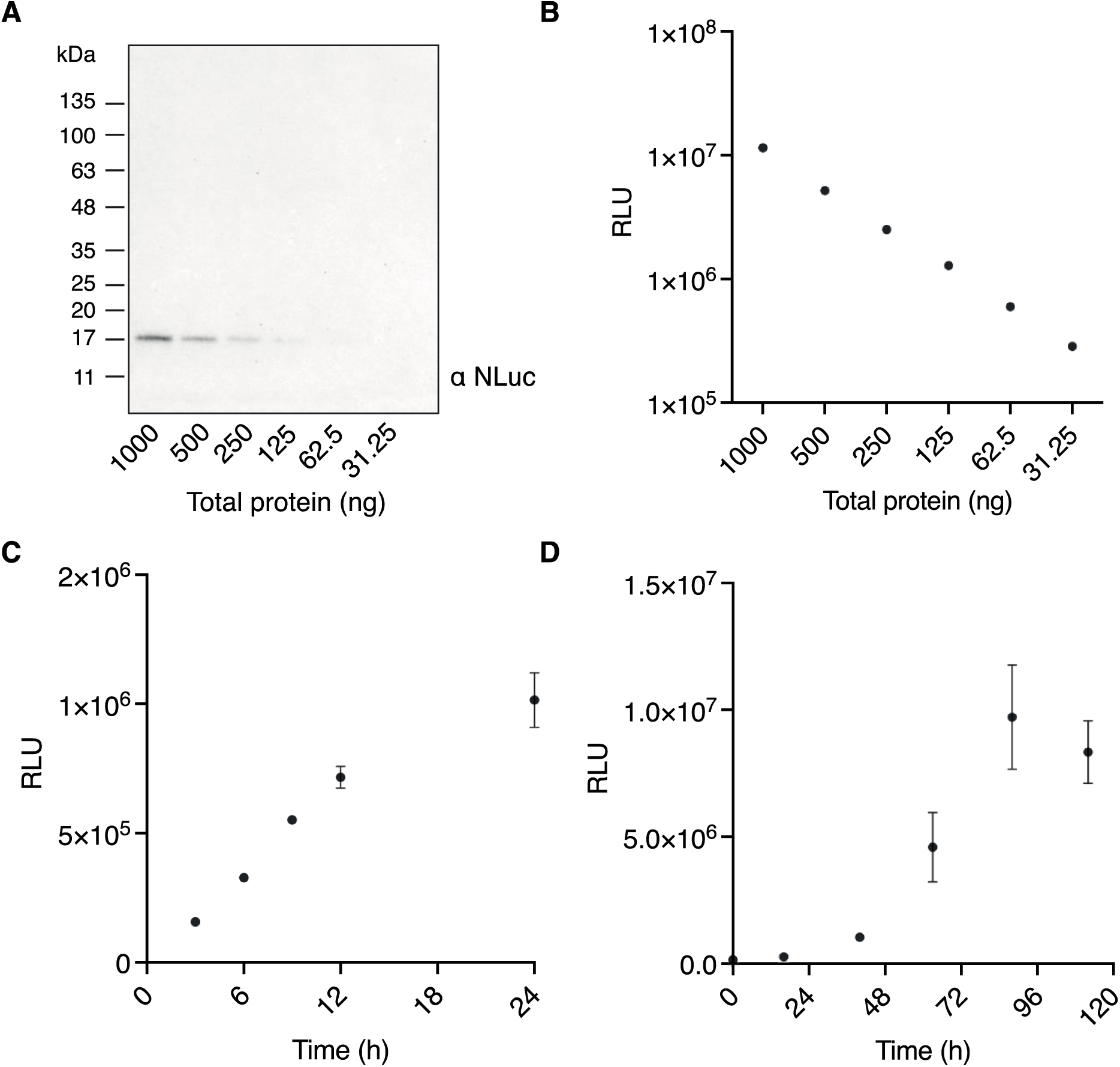
Testing the NLuc secretion reporter. A) Immunodetection of NLuc in a culture medium dilution series ranging from 1 µg to 31.25 ng of total protein. B) Detection of NLuc by luminometry in a culture medium dilution series ranging from 1 µg to 31.25 ng of total protein. C) and D) Time courses of NLuc secretion. Assays were performed with 10 µL of cell-free culture medium, 50 µL of Promega Nano-Glo Assay Buffer and 140 µL of P4 medium. All results are an average of three replicates. Nonvisible error bars are smaller than the data symbol.

*In vivo* secretion time series are particularly challenging due to the low levels of secreted proteins present in the early stages of cell growth. Therefore, we proceeded to test whether the NLuc reporter assay could be used to assess protein secretion dynamics. Mid-exponential *Synechocystis* cultures harboring the NLuc reporter were diluted to an OD_750_ = 0.25 and induced. NLuc secretion was visible already after 3 h of growth and steadily increased over a 24 h period (Fig. 2C). The time series analysis was then extended to a 5-day period (Fig. 2D). NLuc accumulation is visible until 96 h, after which protein levels decrease. Establishing a growth curve by measuring OD was not possible due to the strong aggregating phenotype typical of the *Synechocystis* strain used here (9). Aggregation also led to larger variation in replicate measurements after 48 h. In summary, these results show that the secretion machinery is engaged from the onset of growth and that protein release is gradual and consistent over time.

### T4aP are dispensable for non-pilin protein secretion

Several studies in cyanobacteria have associated changes in the exoproteome to defects in T4aP assembly (22, 23). Therefore, we aimed to investigate whether the T4aPS is involved in the secretion of non-pilin proteins or if the observed changes in the exoproteome resulted from pleiotropic effects. To this end, we conjugated the plasmid pDAR25, containing the NLuc secretion reporter, into a collection of T4aPS mutants (Table 1) to assess the difference in secretion levels between them. T4aPS mutants are known to exhibit diverse aggregation phenotypes (9). During this study, we observed that, in later growth stages, aggregation led to a larger variability in secretion levels (Fig. 2D). Therefore, luminescence was measured after 24 h and normalized to final OD_750 nm_ to minimize aggregation effects.

**Table 1.** List of *Synechocystis* strains used in this study

| Strain | Gene inactivated | Description | Reference |
| --- | --- | --- | --- |
| WT substrain 'PCC-M' | – | – | – |
| $\Delta hfq$ | *ssr3341* | RNA chaperone | (27) |
| $\Delta pilA1$ | *sll1694* | Major pilin | (9) |
| $\Delta pilQ$ | *slr1277* | Outer membrane pore | This study |
| $\Delta pilT1$ | *slr0161* | Pilus retraction ATPase | (9) |
| $\Delta pilA5-6$ | *slr1928, slr1929* | Minor pilins | (12, 28) |
| $\Delta pilA9-slr2019$ | *slr2015, slr2016, slr2017, slr2018, slr2019* | Minor pilins | (9, 12) |
| $\Delta cya1$ | *slr1991* | Adenylyl cyclase | This study |

We started by comparing secretion levels between the *Synechocystis* WT and the Δ*hfq* mutant. The cyanobacterial Hfq protein is a homolog of the RNA chaperone Hfq. It binds to the pilus base via the C-terminus of the extension ATPase PilB1. Δ*hfq* mutants are non-motile, not naturally competent, lack thick pili but retain thin pili (12, 27, 29). Therefore, we hypothesized that, if the T4aPS was involved in the secretion of non-pilin proteins, the Δ*hfq* mutant would show impaired secretion. Accordingly, NLuc was secreted at significantly lower levels in the Δ*hfq* mutant (approximately 27% of the WT) (Fig. 3). The altered secretion levels could also be due to pleiotropic effects since disruption of Hfq is known to alter the transcript accumulation of many genes (27, 29). To further explore this, we analyzed NLuc secretion levels in mutants of the major pilin, PilA1 and the outer membrane secretion pore, PilQ. If the T4aPS was, in fact, mediating non-pilin protein secretion, then a decrease in secretion levels should also be observed when disrupting these core T4aPS components. However, this was not the case. Between the WT and the Δ*pilQ* mutant no significant difference was observed and, remarkably, NLuc secretion levels in the Δ*pilA1* mutant were more than twice as high when compared to the WT. We proceeded to measure NLuc secretion levels in a Δ*pilT1* mutant and two minor pilin mutants (Δ*pilA5-6* and Δ*pilA9-slr2019*). Secretion in the Δ*pilT1* mutant was significantly reduced to a level close to that observed in the Δ*hfq* mutant. In the minor pilin mutants, a small, but significant decrease was observed. Taken together, these results show that T4aP are dispensable for non-pilin protein secretion. However, altered secretion efficiency in certain pili mutants suggests an indirect role of the T4aPS on the secretion of non-pilin proteins.

**Figure 3.**
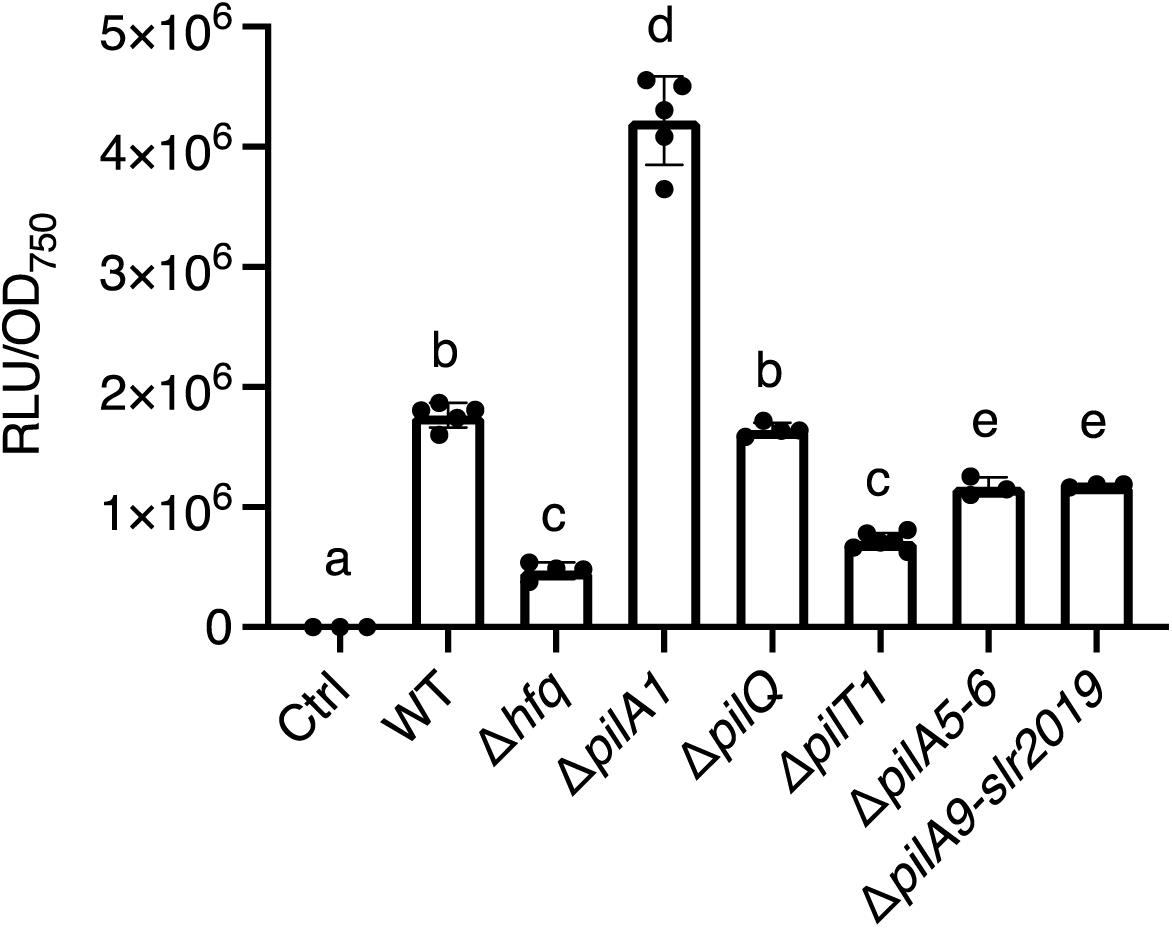
NLuc secretion assays. Assays were performed with 10 µL of cell-free culture medium, 50 µL of Promega Nano-Glo Assay Buffer and 140 µL of P4 medium. Results are an average of three to six replicates and full circles represent individual replicate values. Significance letters were attributed by comparing mean luminescence values using a one-way ANOVA followed by a Tukey’s multiple comparisons test. Ctrl designates a pDF-trc empty vector control strain.

### Different secretion levels are consistent with differences in NLuc accumulation at the cytoplasmic membrane

The observation that Hfq and PilT1, but not PilA1 or PilQ, was required for efficient NLuc secretion was surprising. Therefore, we aimed to investigate the reasons behind the repression of secretion. In *Synechocystis*, the second messenger 3′, 5′-cyclic AMP (cAMP) is involved in the regulation of surface features of the cell and many cAMP-dependent transcripts were affected in the Δ*hfq* mutant (27, 30). Therefore, we proceeded to determine whether the deregulation of cAMP signaling was altering secretion profiles. For this experiment, we first tested a Δ*cya1* mutant harboring the NLuc reporter. Cya1 is an adenylate cyclase responsible for producing virtually all intracellular cAMP in *Synechocystis* (31). The Δ*cya1* mutant is non-motile, but motility can be recovered by the addition of extracellular cAMP (32). Therefore, we hypothesized that, if cAMP is involved in the regulation of protein secretion, the Δ*cya1* mutant would show impaired NLuc secretion which could be recovered by the addition of extracellular cAMP. This was not the case, as the Δ*cya1* mutant presented WT levels of NLuc secretion and no significant difference was observed in the presence of extracellular cAMP (Fig 4A). NLuc secretion levels were also measured in the remaining *Synechocystis* mutants in the presence of cAMP. When compared to secretion levels in the absence of cAMP (Fig. 3), no significant difference was observed (Fig. 4B). The data allow us to conclude that NLuc secretion is not dependent on the intracellular levels of cAMP.

**Figure 4.**
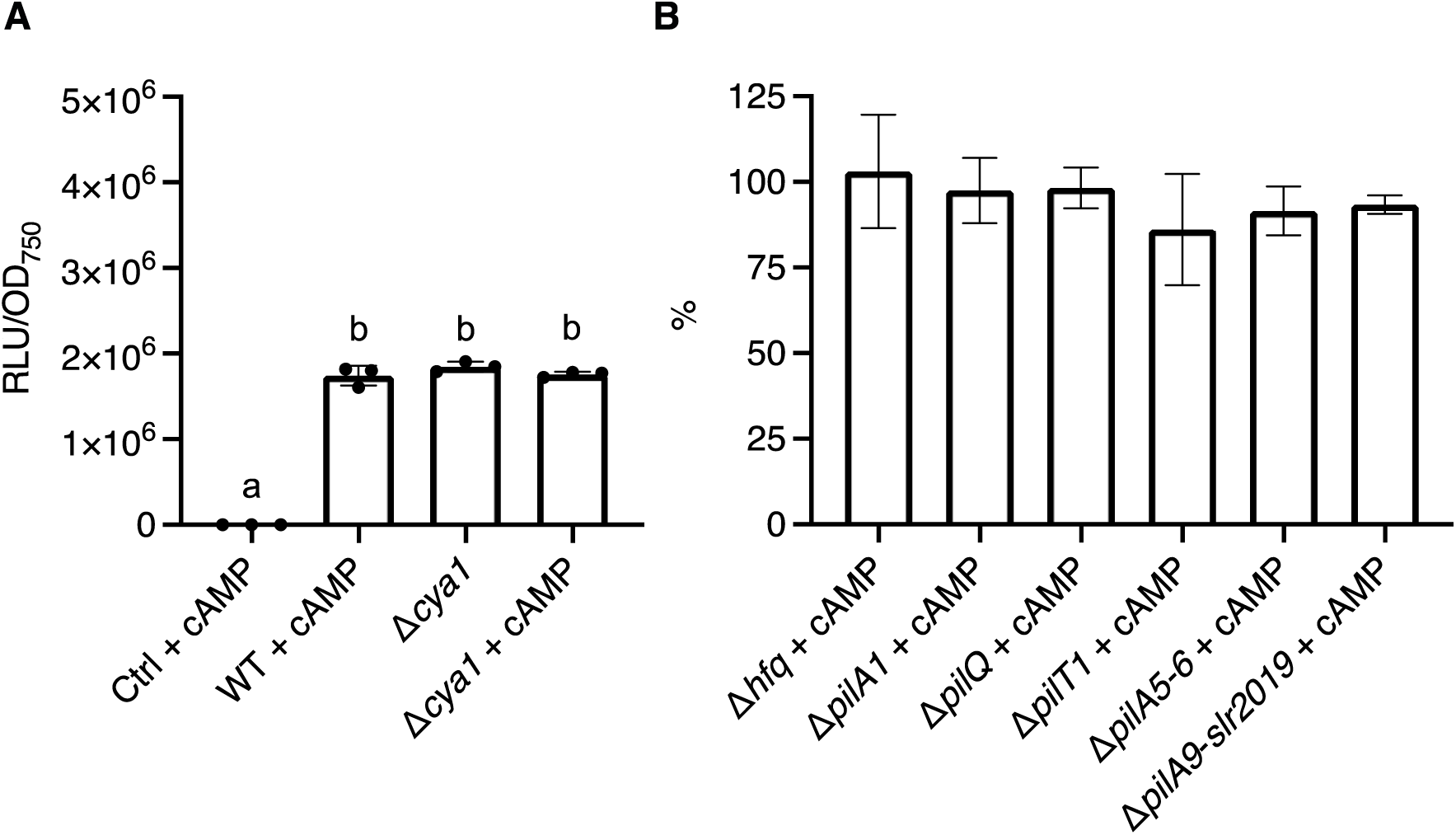
NLuc secretion assays in the presence of cAMP. A) Normalized luminescence values for the WT, the Δ*cya1* mutant and the empty vector control strain (Ctrl). B) Normalized luminescence values of the T4aPS mutants in the presence of cAMP as a percentage of the values in the absence of cAMP (see Fig. 3). When indicated, 100 µM cAMP was added at the point of induction. Results are an average of three to four replicates and full circles represent individual replicate values. Significance letters were attributed by comparing mean luminescence values using a one-way ANOVA followed by a Tukey’s multiple comparisons test.

Having ruled out any effect of cAMP on NLuc secretion levels, we investigated whether disruption of the T4aPS could lead to cell envelope stress, and consequentially, a decrease in secreted NLuc. In our previous study, we observed that lower secretion levels with a Tat signal peptide were caused by inefficient translocation of the target protein across the cytoplasmic and outer membranes. This resulted in visible protein degradation and accumulation of the protein in the cytoplasmic membrane and periplasm (24). Therefore, the *Synechocystis* mutants were fractionated into spheroplasts (containing thylakoid membrane, cytoplasm and cytoplasmic membrane) and periplasm to see if a similar phenomenon could be observed. Strikingly, the spheroplast fractions of the secretion impaired *Δhfq* and *ΔpilT1* mutants showed visible signs of protein degradation and strong accumulation of both the pre-protein and mature NLuc. On the other hand, in the secretion enhanced *ΔpilA1* mutant, neither protein accumulation nor degradation was observed. Accumulation of mature NLuc is visible in the periplasm fraction of all tested strains, however, no clear trend can be observed.

## Discussion

In this study, we have established a quantitative, luminescence-based reporter to study two-step protein secretion in cyanobacteria. The reporter was generated by fusing the *Tf*AA10A Sec signal peptide, previously shown to mediate secretion in cyanobacteria (24), to NLuc. As a proof of concept, the reporter was used to determine if the T4aPS fulfills a dual role of secreting both pili and non-pilin proteins. Results show that the assembly of T4aP is not required for NLuc secretion. Moreover, secretion impairment in T4aP-deficient mutants is likely due to perturbation of cell envelope homeostasis rather than the absence of T4aP.

### NLuc is a superior reporter of protein secretion in cyanobacteria

The elucidation of protein secretion mechanisms in cyanobacteria has been impeded by the lack of assays to quantitatively follow secretion dynamics. Here, we take a first step to tackle this bottleneck by developing a sensitive NLuc-based secretion reporter. NLuc secretion could be detected with small volumes, short handling time and across a wide dynamic range. Therefore, secretion dynamics could be monitored from the onset of growth and followed throughout the entire growth curve with negligible impact on culture volume. Common options to monitor protein secretion in Gram-negative bacteria include *Escherichia coli* proteins with extracytoplasmic location (e.g. alkaline phosphatase PhoA and TEM β-lactamase BlaM) and a variety of fluorescent proteins (33). However, both present disadvantages. PhoA and BlaM require disulfide bond formation, therefore protein folding in the cytoplasm, a requirement for Tat-targeted proteins, is not possible (34). On the other hand, fluorescent proteins, such as GFP and variants thereof, can only be secreted after cytoplasmic folding. As a result, they are not compatible with translocation through the Sec pathway (35, 36). NLuc does not suffer from these shortcomings. In this study we have shown that NLuc can fold outside the cytoplasm and that it can be used to monitor two-step secretion with a Sec signal peptide. Previous studies have shown that NLuc is also active in the cyanobacterial cytoplasm (37, 38), suggesting compatibility with Tat-mediated protein secretion. In addition, NLuc has recently been used as a reporter for the one-step type III secretion systems of *Salmonella* and *Yersinia* (39, 40), thus it is likely also compatible with cyanobacterial one-step secretion systems such as the T1SS. In sum, NLuc is a robust and versatile reporter with the potential to facilitate future protein export and secretion studies in cyanobacteria.

### Core elements of the T4aPS are not required for NLuc secretion

Having developed the NLuc secretion reporter, we aimed to answer whether the cyanobacterial T4aPS can mediate two-step secretion of non-pilin proteins. The proposed dual role of the T4aPS in pili assembly and non-pilin protein secretion is supported by the literature. First, in several species of Proteobacteria the type IV pili system been shown to secrete both pilins and non-pilin proteins (16–18, 41). This has most recently been observed in *Geobacter sulfurreducens* where pili were shown to be essential for the secretion of OmcS and OmcZ nanowires (19). Second, in cyanobacteria, there are several studies where pilin signal peptides have been used to mediate heterologous protein secretion (42–44). Finally, a series of studies have shown that inactivation of T4aPS proteins, such as PilB1 and Hfq, altered both the exometabolome and the exoproteome (22, 23, 45, 46). Despite this body of work supporting the hypothesized dual role of the T4aPS, our results suggest otherwise.

In line with these studies, NLuc secretion was significantly decreased in the Δ*hfq* background (Fig. 3). However, since inactivation of Hfq is known to have a pleiotropic effect on the cell (27, 29), we proceeded with a more targeted analysis. Surprisingly, contrary to the initial hypothesis, inactivation of key T4aPS components, such as the major pilin PilA1 and the outer membrane pore PilQ, did not negatively affect NLuc secretion (Fig. 3). In the Δ*pilQ* background, NLuc secretion levels were similar to the WT. In *Francisella novicida*, deletion of PilQ resulted in a significant reduction of protein secretion (17) and in *Shewanella*, GspD, the PilQ-like secretin of the T2SS, is essential for the secretion of T2SS substrates (47). Therefore, this is a strong indication that the T4aPS is not directly involved in the secretion of non-pilin proteins in cyanobacteria.

Interestingly, NLuc secretion in the Δ*pilA1* background significantly increased (Fig. 3). PilA1 is one of the major proteins transported across the cytoplasmic membrane and accumulation of unprocessed PilA1 prepilin leads to the degradation of the Sec-YidC complex (48). Therefore, the enhancement of secretion in the absence of PilA1 is likely due to an increased capacity of the Sec pathway. This hypothesis was corroborated by our location study where neither NLuc degradation nor accumulation in the spheroplasts was observed (Fig. 5). Going forward, the deletion of native proteins competing for the Sec and Tat pathways may be of interest for biotechnological applications as a strategy to increase heterologous protein export and secretion.

**Figure 5.**
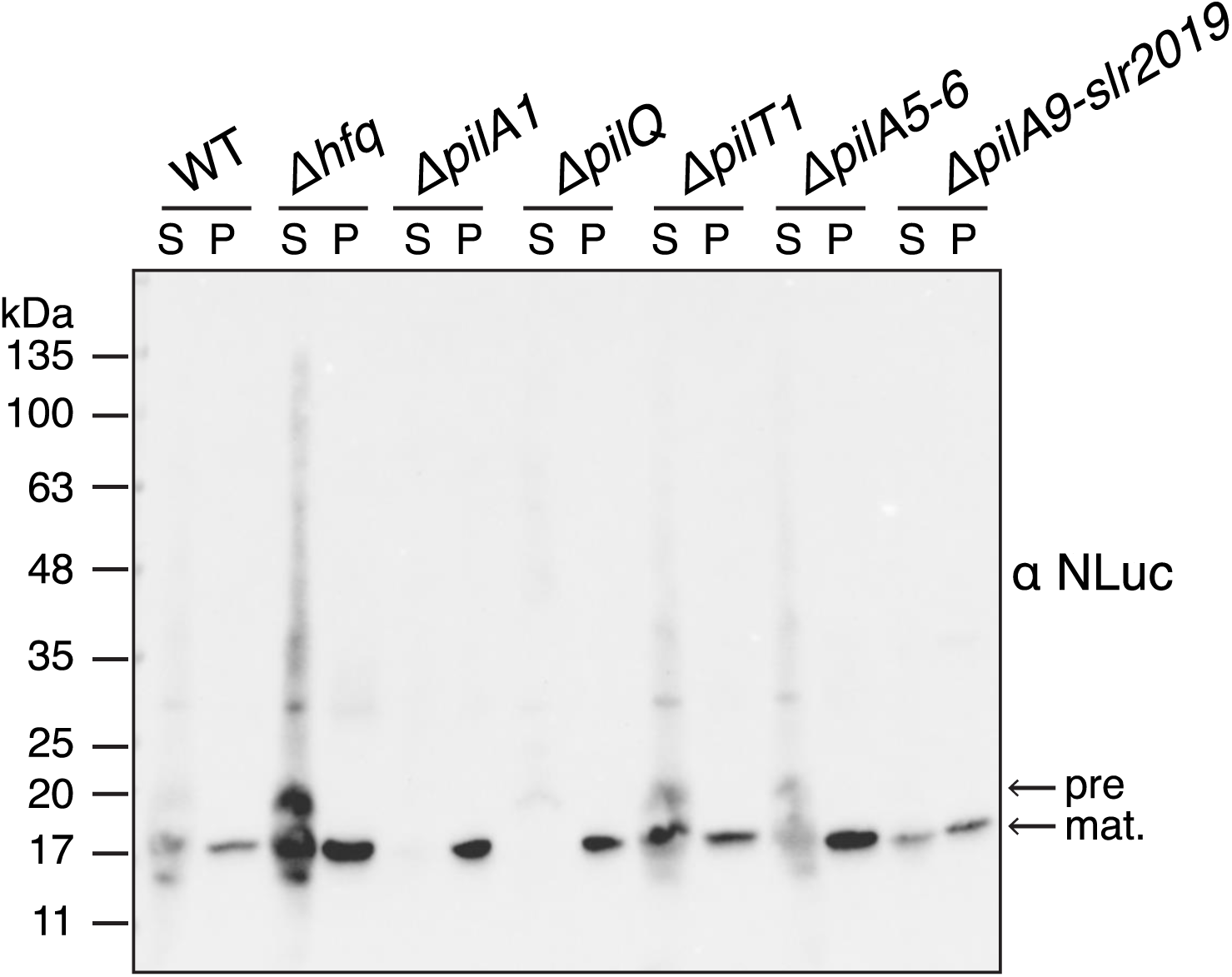
Location study of NLuc in *Synechocystis* mutants. NLuc was detected in spheroplast (S) and periplasm (P) fractions by immunoblot analysis with an NLuc antibody (αNLuc). Lanes were normalized to an equivalent of OD_750 nm_ = 0.25. Pre- and mature protein bands are indicated with arrows.

### Envelope stress responses may explain secretion phenotypes in T4aPS mutants

Our first hypothesis to explain the differing NLuc secretion levels was the disruption of cAMP signaling (27), however the addition of cAMP did not alter secretion levels in any of the tested strains (Fig. 4A, B). It is worth highlighting that, in our analysis, a non-motile *Δcya1* mutant, which is deficient in cAMP production (31), showed WT NLuc secretion levels both with and without the addition of exogenous cAMP. Therefore, it was concluded that the intracellular levels of cAMP do not influence protein secretion. An alternative hypothesis, that is supported by the NLuc location study (Fig. 5), is that the disruption of the T4aPS may alter the correct functioning of the cell envelope. It is well-documented that changes to the T4aPS can lead to altered cell surface properties and cell morphology (7, 48). However, little is known regarding how elements of the T4aPS can regulate, directly or indirectly, cytoplasmic and outer membrane protein composition. In *E. coli*, Hfq is an RNA-binding chaperone that localizes to the cytoplasmic membrane (49) and, through several small RNAs (sRNAs), controls the expression of outer membrane proteins at the post-transcriptional level (50). In *Synechocystis* and *S. elongatus*, Hfq localizes to the cytoplasmic membrane and forms a complex with PilB1 which is essential for T4aP assembly (23, 29). Although a role in RNA binding has not yet been demonstrated, the inactivation of Hfq leads to reduced accumulation of several mRNAs and sRNAs which include transcripts of several outer membrane proteins (27, 29). Specifically, the two most downregulated genes in the Δ*hfq* mutant, *cccS* and *cccP*, may be involved in the construction of cell surface components (51). Therefore it is conceivable that Hfq, directly or indirectly, influences the composition of the cyanobacterial cell envelope.

Secretion defects can also derive from increased competition for access to the Sec translocon. Our results showed that NLuc secretion levels were significantly lower in a *ΔpilT1* mutant (Fig. 3). In *Synechocystis*, disruption of *pilT1* leads to a five to seven times increase in the accumulation of *pilA1* mRNA (7). Together with the observation that NLuc secretion levels increased in a Δ*pilA1* mutant, we can hypothesize that an increased pool of PilA1 prepilin may jam the Sec machinery and cause the NLuc accumulation visible in the spheroplasts of the Δ*pilT1* mutant (Fig. 5).

The Gram-negative cell envelope is a complex system with a myriad of interdependent components. The emerging picture shows that the T4aPS is tightly linked with the cell envelope network and perturbation of pili biogenesis and assembly leads to pleiotropic phenotypes. A compromised cell envelope could also explain the many reported defects in compound uptake and secretion in cyanobacterial T4aPS mutants and substrains with impaired T4aP biogenesis (52, 53). However, the routes of protein transport across the outer membrane in two-step secretion in cyanobacteria remain unclear. The predominant outer membrane proteins of *Synechocystis* allow only the passage of inorganic ions (54). Therefore, one possibility is the existence of a less abundant outer membrane pore which is permeable to organic solutes. Non-classical secretion routes, such as outer membrane vesicles, could also play a role (6). In conclusion, this study provides novel insights into protein secretion in cyanobacteria, and we expect that the secretion reporter developed here will contribute to further advances in the field.

## Materials and methods

### Strains and growth conditions

The motile *Synechocystis* sp. PCC 6803 substrain ‘PCC-M’, originally obtained from S. Shestakov (Moscow State University, Russia), was used in this study. All cultures were maintained on BG-11 medium (55) supplemented with 10 mM 2-tris(hydroxymethyl)-methyl-2-amino 1-ethanesulfonic acid (TES) buffer (pH 8.0) and 1.5% (w/v) bacto agar at 30°C with continuous illumination of approximately 25 μmol photons m^−2^ s^−1^. Liquid cultures were grown in P4-TES CPH medium (24), a modified version of phosphate replete medium (56), in a two-tier vessel system (HDC 6.10 starter kit CellDEG, Germany) where CO_2_ is supplied to the cultures through a permeable polypropylene membrane (57). The lower tier contained 200 mL of carbonate buffer obtained by combining solutions of 3 M KHCO_3_ and 3 M K_2_CO_3_ at a ratio of 4:1 yielding a CO_2_ partial pressure of 32 mbar (reference T = 20°C). Into the upper tier, 25 mL growth vessels with 10 mL of growth medium were inserted. The system was illuminated by LUMILUX cool white L 15W/840 fluorescent lamps (Osram, Germany) with continuous illumination of approximately 40 μmol photons m^−2^ s^−1^ and shaken at 280 rpm on a Unimax 1010 orbital shaker (Heidolph Instruments, Germany). Agar plates were supplemented with the respective antibiotics (25 μg mL^−1^ spectinomycin, 25 μg mL^−1^ streptomycin, 25 μg mL^−1^ apramycin, 25 μg mL^−1^ kanamycin, 15 μg mL^−1^ chloramphenicol). During experiments, liquid cultures were grown without antibiotics to prevent distorting effects. If not stated otherwise, cultures were induced with 100 µM isopropyl β-d-1-thiogalactopyranoside (IPTG) for 24 h before measurements.

### Generation of Synechocystis *mutants*

*Hfq, pilA1, pilA5-6, pilA9-slr2019* and *pilT1* mutants were generated in previous studies (Table 1). The *pilQ* deletion mutant was generated by transforming *Synechocystis* with chromosomal DNA of a *pilQ* mutant from a non-motile substrain (48). Transformants were streaked on BG-11 agar plates with increasing concentrations of chloramphenicol up to 7 µg ml^-1^ and full segregation was tested by colony PCR. The *cya1* deletion mutant was generated by replacement of the native locus with a spectinomycin resistance gene via homologous recombination. The flanking regions were amplified from the *Synechocystis* genome through PCR and assembled with an *aadA* resistance cassette into a pRL271-based vector backbone by Gibson assembly using NEBuilder HiFi DNA Assembly Master Mix (New England Biolabs, USA). Transformants were streaked on BG-11 agar plates with increasing concentrations of spectinomycin up to 25 µg ml^-1^ and full segregation was tested by colony PCR. The NLuc secretion reporter was generated by fusing the native Sec signal peptide (AA 1–36) of *Tf*AA10A (Uniprot KB: Q47QG3) (24) to the N-terminus of the full-length amino acid sequence of NLuc (GenBank: JQ437370.1). The construct was custom synthesized by GenScript (USA) with EcoRI and HindIII recognition sequences at the 5’ and 3’ end, respectively. Using the EcoRI and HindIII recognition sequences, the construct was inserted into the pDF-trc plasmid (24, 26) by restriction digest cloning. The sequence of the generated plasmid was confirmed by Sanger sequencing and named pDAR25. Due to the incompatibility of the pDAR25 *aadA* antibiotic resistance gene with the *cya1* deletion mutant, a second version of pDAR25 was generated by replacing the *aadA* resistance cassette with *aph(3’)-Ia*, which confers resistance to kanamycin, by Gibson assembly using NEBuilder HiFi DNA Assembly Master Mix (New England Biolabs, USA). This vector was named pDAR25-Kan. pDAR25 and pDAR25-Kan were introduced into *Synechocystis* by biparental mating as previously described (24) with the exception that spectinomycin or kanamycin were used at a concentration of 25 μg mL^−1^ and selection plates for the T4aPS mutants included their respective background antibiotic.

### NanoLuc luciferase assays

To measure NLuc activity in the extracellular space, culture supernatants were separated by centrifugation for 5 min at 10 000 × g. If not stated otherwise, 10 µL of the cleared supernatant was mixed with 50 µL of Promega Nano-Glo Assay Buffer (containing Nano-Glo Assay Substrate at 1:50 ratio) and 140 µL of P4-TES CPH medium (total of 200 µL assay volume). In blank reactions, the cleared supernatant was replaced by P4-TES CPH medium. All assay components were allowed to equilibrate to room temperature before addition. Following addition, the assay was incubated at room temperature for 3 min before measurement. Luciferase activity was measured in Nunc black, optical bottom, 96-well plates (Thermo Fisher Scientific, Germany) on a Thermo Varioskan Flash microplate reader (Thermo Fisher Scientific, Germany) with the dynamic range set to autorange and measurement time of 10000 ms. For the NLuc time series, a mid-exponential culture of *Synechocystis* harboring the NLuc reporter was diluted to an OD_750 nm_ = 0.25 and induced with 100 µM IPTG. Samples of cell free culture medium were collected every 3 h and measured immediately after processing. For comparison of NLuc secretion in the T4aPS mutants, mid-exponential cultures were diluted to an OD_750 nm_ = 0.25 and induced with 100 µM IPTG. The cAMP addition assays included 100 µM cAMP at the point of induction. Samples were harvested at 0 and 24 h and measured immediately after processing. Luminescence values at 24 h were subtracted of time zero and normalized by OD_750 nm_ of the culture. Comparison of the normalized luminescence values was done with a one-way ANOVA followed by a Tukey’s multiple comparisons test performed in Prism (version 9.0, GraphPad Software, USA).

### Preparation of cellular and extracellular fractions for immunoblotting

Cell lysates were prepared by centrifuging cultures 5 min at 10 000 × g and resuspending in an appropriate volume of lysis buffer (20 mM Tris–HCl pH 7.5, 1% SDS, 10 mM NaCl). Cells were broken with zirconium oxide beads (diameter: 0.15 mm) using a Bullet Blender Storm 24 (Next Advance, USA) with the following settings: 3 times 5 min intervals at level 12. Lysates were centrifuged at 10000 × g for 10 min at 4°C to remove cell debris and transferred to fresh tubes for protein content determination. Culture supernatants were prepared by centrifuging cultures 5 min at 10000 × g and concentrating the cell free supernatant approximately 10 times with 3 kDa molecular weight cut-off Amicon Ultra-15 centrifugal concentrators (Merck Millipore, Germany). The concentrated supernatant was then transferred to fresh tubes for protein content determination. Periplasmic fractions were extracted by cold osmotic shock following a previously described protocol (58) from culture volumes equivalent to OD_750 nm_ = 5. The resulting periplasm and spheroplast fractions were lyophilized, resuspended in 100 µL of sample loading buffer (50 mM Tris-HCl pH 6.8, 2% SDS, 0.1% bromophenol blue, 10% glycerol and 100 mM dithiothreitol (DTT)) and incubated for 5 min at 95°C prior to separation by SDS-PAGE. For the NLuc location study, 5 µL of each fraction were separated by SDS-PAGE.

### SDS-PAGE and immunoblot analysis

Protein content of samples for SDS-PAGE and immunoblotting was analyzed using a Pierce BCA Protein Assay (Thermo Scientific, Germany) with a standard curve from 0 to 2000 µg mL^−1^ bovine serum albumin following the manufacturer’s recommendations. Samples were incubated for 5 min at 95 °C with sample loading buffer prior to separation by SDS-PAGE. Proteins were separated on SERVA Neutral HSE gels (Serva, Germany) in Laemmli running buffer (25 mM Tris, 192 mM glycine and 0.1% SDS) at 300 V. After SDS-PAGE separation, proteins were transferred onto a 0.45 µm nitrocellulose membrane, using a BlueBlot Semi-Dry Blotter (Serva, Germany), for 30 min at 12 V. Membranes were blocked with 5% skimmed milk powder (w/v) in TBS-T buffer (10 mM Tris-HCl pH 8.0, 150 mM NaCl and 0.05% Tween-20) for 60 min at room temperature. Subsequently, the membranes were incubated in anti-NLuc 1:5000 (R&D Systems, USA) or anti-RbcL 1:5000 (large subunit, form I and II, AS03 037) (Agrisera, Sweden) at 4°C overnight. Membranes were washed with TBS-T and incubated for 60 min at room temperature with an anti-mouse or anti-rabbit horseradish peroxidase (HRP)-conjugated secondary antibody (Promega, Germany) at dilutions of 1:5000. Membranes were washed again with TBS-T and the HRP signal was developed using Pierce ECL Western Blotting Substrate (Thermo Fisher Scientific, Germany) and detected with a ChemiDoc imaging system (Bio-Rad Laboratories, Germany).

## Data availability

Vector maps and sequences for pDAR25 and pDAR25-Kan can be found in Supplemental Material S1 at Zenodo doi:10.5281/zenodo.5541267.

## Acknowledgements

D.A.R. was supported by the Alexander von Humboldt Foundation. Work in the laboratory of P.E.J. was supported by the Novo Nordisk Foundation (NNF16OC0021832 and NNF19OC0057634). C.W.M. and F.D.C were supported by the Leverhulme Trust RPG-2020-054. N.S. and A.W. were supported by the German Science Foundation (WI2014/7-1 and in frame of the SFB1381 - 403222702-SFB 1381 (A2)).

## Conceptualization

D.A.R., J.A.Z.; Methodology: D.A.R.; Investigation: D.A.R., J.A.Z., F.D.C., N.S.; Resources: D.A.R., J.A.Z., F.D.C., N.S., P.E.J., C.W.M., A.W. G.P.; Data curation: D.A.R.; Formal analysis: D.A.R.; Visualization: D.A.R., J.A.Z.; Writing – original draft: D.A.R., J.A.Z.; Writing – review & editing: D.A.R., J.A.Z., F.D.C., N.S., P.E.J., C.W.M., A.W., G.P.; Funding acquisition: D.A.R., P.E.J., C.W.M., A.W.

We declare that we have no conflicts of interest.

